# No Evidence for Communication in the Complete Locked-in State

**DOI:** 10.1101/287631

**Authors:** Martin Spüler

## Abstract

Enabling communication for patients in the complete locked-in state (CLIS) has been a major goal for Brain-Computer Interface (BCI) research over the past 20 years. Last year, two papers were published that claim to have reached this goal: (Chaudhary et al., 2017) were the first to report communication in CLIS using a method based on fNIRS. Few month later, (Guger et al., 2017) claimed that their EEG-based BCI system is able to restore communication in CLIS. This manuscript demonstrates methodological flaws in the analysis of both papers and that their conclusions are invalid. Further, the data from (Chaudhary et al., 2017) is reanalyzed to demonstrate that their results cannot be reproduced and that there is currently no scientifically sound evidence that demonstrates communication in CLIS.

## Article

When Birbaumer and colleagues [1] showed in 1999 for the first time that a person in locked-in state can use a Brain-Computer Interface (BCI) to communicate, it also created the hope of BCIs restoring communication in the complete locked-in state (CLIS), where a patient has no remaining muscle control. Since this pioneering work, multiple Electroencephalography(EEG)-based BCI systems were successfully tested with locked-in patients [2]. However, these systems did not work for complete locked-in patients leading to the conclusion that voluntary brain regulation is not possible in CLIS [2,3].

This changed in 2014, when Gallegos-Ayala et al. [4] presented a case-study suggesting that near-infrared spectroscopy (NIRS) could be used for communication in the complete locked-in state. It was followed up by Chaudhary et al. in 2017 [5], who recorded NIRS in 4 patients in complete locked-in state. In that work, results from offline and online classification are presented with accuracies significantly above chance level, which led the authors to the conclusion that NIRS-based BCI communication is working in CLIS.

For this commentary, i performed a reanalysis of the data from Chaudhary and colleagues [5]. As the results are substantially different from the results reported in the original paper, I question the claim of NIRS-based BCI communication in the complete locked-in state.

## Reanalysis of NIRS data

For the reanalysis, the data were used that were published as supplementary material [6-9] to the original paper. This data contains the preprocessed NIRS signal (HbO) for 20 channels and all trials of all training and feedback sessions. (with the exception of patient F for whom 22 sessions are missing in the data), separated into “true/yes” and “false/no” trials.

### Statistics

Two different statistical analyses were performed and both showed no significant difference in the NIRS response between yes/no questions. The first analysis was different to the one performed by Chaudhary et al. and is presented in the appendix in more detail. For the second analysis I point out methodological flaws in the statistical analysis of Chaudhary et al. and show results with corrected methods.

As the statistical analysis by Chaudhary et al. was not described with sufficient detail to exactly reproduce it, Chaudhary and colleagues responded in the first review round to this comment, that they first averaged the data over all trials, and then over all sessions and performed a t-test on those averages. The problem with this kind of analysis is that the variance over trials/sessions is removed by the averaging and only the variance over the channels is retained. Performing a statistical test will then compare the mean of yes-trials with the mean of no-trials, while considering the variance over all channels. As the channels are highly correlated (not independent), the variance is very low and will lead to the wrong result that the difference is significant. That this kind of analysis is not correct can be tested by using a permutation test. Using a random permutation of the trials, instead of a separation by yes/no, should show no significant difference if the statistical analysis is correct. But for the method used by Chaudhary and colleagues, the results are significant in each of ten random permutations (see figure A1 in appendix), which demonstrates that the used method is not correct.

This does not mean that averages should not be used for statistical analysis, but that the order of averaging matters. If the data is first averaged over channels, then over trials, the variance over sessions is retained. But in this case, there is no significant difference between yes/no questions (*p* > 0.05; t-test, not corrected for multiple comparisons) for any patient. Figure 1 shows the results of an analysis of data from patient B with the two different orders of averaging. While the mean response is the same for both analyses, the variance and the results of a statistical test are different.

**Fig 1.**
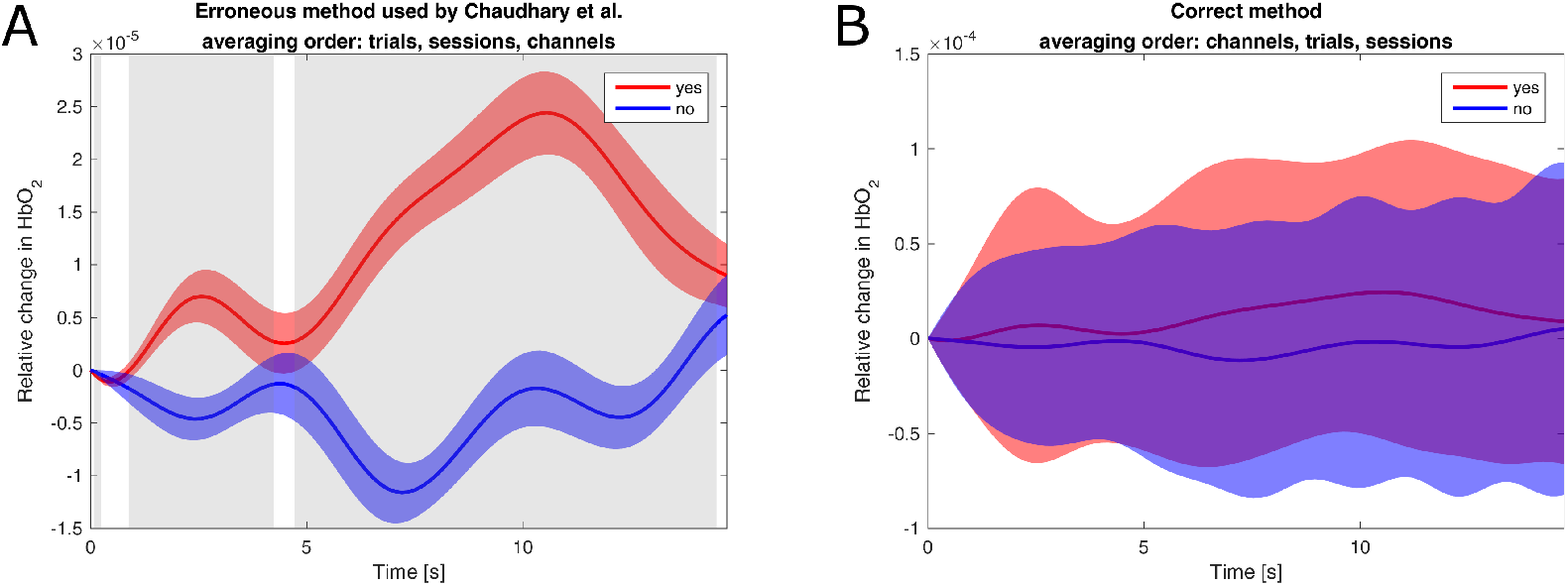
Comparison of statistical methods on data of patient B. Both plots show the average response to yes/no questions as well as the standard deviation. A: Method used by Chaudhary et al. First averaging over trials, then over sessions, then over channels. For the t-test the variance over channels is retained and used. Timepoints with significant difference (*p* < 0.0005) are marked grey. B: Same method, but with a correct order of averaging. First averaging over channels, then over trials, then over sessions. For the t-test the variance over sessions is retained and used. No timepoints show a significant difference (*p* > 0.05).

### Offline classification

The offline classification of the data was also reproduced. As some details of this analysis were not given by Chaudhary and colleagues (e.g. hyperparameter) it was not possible to reproduce the results with exactly the same method, but a similar approach was used. A support vector machine (SVM) with a linear kernel was applied to the relative change in HbO_2_. For each day of each patient, a 10-fold cross-validation was performed in which the data was randomly divided into 10 blocks and 9 blocks were used for training the classifier and tested on the remaining block. This process was repeated 10 times, so that each block was used for testing once. For training the classifier, the training data was balanced by randomly removing trials of the majority class from the training data. The optimal hyperparameter *C* for the SVM was estimated by performing a gridsearch with a 10-fold crossvalidation on the training data. When using the preprocessed data as input for classification, an average accuracy of 49.4 % was obtained. The performance was not significantly different [10] than chance level (*p* > 0:05, not corrected for multiple comparisons) for all of the 42 days. More detailed results can be found in the appendix.

## Discussion

In summary, a reanalysis of the data from Chaudhary et al. [5] has shown no significant difference in the hemodynamic response to “yes” and “no” questions, and the NIRS data could not be classified with an accuracy significantly above chance level.

As the obtained results are in stark contrast to the results presented by Chaudhary et al., possible reasons should be discussed. For the statistical part, it was possible to pinpoint the methodological flaw and therefore explain the erroneous results. Due to the lack of details in the description of the offline classification of Chaudhary et al. (e.g. choice of hyperparameter), one can only speculate and it is up to Chaudhary and colleagues to provide more information and an explanation why their results can not be reproduced and why they report online accuracies above 70 %.

As the data recorded by Chaudhary and colleagues shows that their approach of a NIRS-based BCI does not work, it should be discussed if the current literature shows any evidence for communication in CLIS. The work of Gallegos-Ayala et al. [4] is from the same research group as Chaudhary et al., uses the same approach and the same patient (Pat. F), therefore can also be seen as disproven.

Regarding the use of EEG-based BCIs for communication in CLIS, Guger et al. [11] have recently claimed that their mindBEAGLE system, using a P300 paradigm, works also for CLIS patients. However, Guger and colleagues rest that claim on a very small sample size of 10 trials and do not provide any statistical assessment of their results. When performing a statistical analysis on that data, the classification accuracies are actually not significantly (*p* > 0.05) above chance level (see appendix for details).

Establishing communication in the complete locked-in state is a major goal for non-invasive BCI research that has been worked on for nearly two decades without success. The recent claims, that this goal has been reached, rest in the case of Chaudhary et al. on a flawed method and erroneous results, or in the case of Guger et al. are just claims without any statistical support. Unfortunately, it has to be concluded that there currently is no scientifically sound evidence that demonstrates communication in the complete locked-in state.

## Supporting information

S2 Matlab Scripts used for analysis

## Supporting information

**S1 Appendix. Appendix** containing more detailed description of the methods and results of the reanalysis.

**S2 File. Zip-Archive with scripts** contains all Matlab scripts that were used for the statistical analysis, offline classification and generating figures for this comment.

## Acknowledgments

This work is supported by the *Deutsche Forschungsgemeinschaft* (DFG; grant SP-1533\2-1).

