## Supplementary material for "No Evidence for Communication in the Complete Locked-in State": S1 Appendix

Martin Spüler

### A1. Statistical analysis originally performed for this comment

In the following, the statistical analysis is described that was originally performed for the comment, but was replaced with a different analysis after the first review round of the comment.
For each patient separately, the data of all sessions were used and a Wilcoxon's ranksum test was performed for each channel and each sample. The minimum p-value for Patient F was $p=0.0026$, for Patient G $p>0.05$, for Patient B $p=0.0317$, and for Patient W $p=0.0109$.
Due to the large number of tests involved (20 channels with 93 samples equals 1860 tests), the results must be corrected for multiple comparisons. Using a Bonferroni correction with a significance level of $\alpha=0.05$, the adjusted significance level is $\alpha / 1860 = 0.000027$, showing that there is no significant difference in the hemodynamic response between "yes" and "no" questions for any of the 4 patients.

### A2. Results from analysis of Chaudhary et al. with randomly permuted trials

In the first review round, Chaudhary and colleagues responded to this comment and gave a detailed description of the statistical procedure performed. They averaged the data first over all trials of one session, then over all sessions. By using this order of averaging, the variance over trials/sessions is removed, retaining only the variance over channels, which is low as channels are not independent and highly correlated. Therefore a statistical analysis will show (erroneously) significant results.
That this method is not correct can be easily shown by performing a permutation test, in which the yes/no labels of all trials are randomly permuted. In a permuted dataset, a correctly applied statistical analysis should not show any significant effects.
However, using the (incorrect) method of Chaudhary et al. shows significant effects as shown in figure A1.

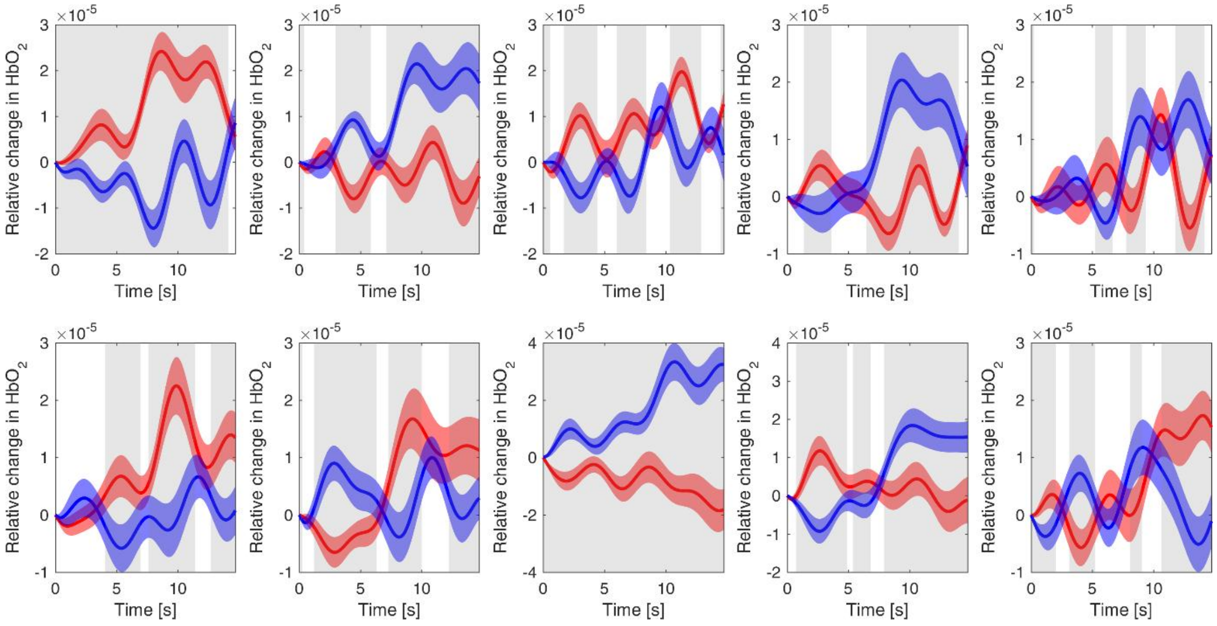


*Figure A1. Method used by Chaudhary and colleagues applied to the data of Patient B with randomly permuted trials. For each subplot, the same data was used, but with a different permutation. Timepoints with a significant difference ($p<0.0005$, t-test) are marked in gray. Red and blue line show the average of two groups of randomly chosen trials, with the areas indicating the standard deviation*

### A3. Classification results

**Table A1.** Day-wise cross-validation results for Patient F with Accuracy (ACC) and Chance level (CL) for Alpha=0.05; No days have a performance significantly (p>0.05) above chance level (not corrected for multiple comparisons).

| Day | D1 | D2 | D3 | D4 | D5 | D6 |
|---|---|---|---|---|---|---|
| Acc | 0.484 | 0.5325 | 0.5275 | 0.4963 | 0.489 | 0.445 |
| CL | 0.5804 | 0.5895 | 0.5736 | 0.5895 | 0.5804 | 0.5895 |

**Table A2.** Day-wise cross-validation results for Patient G with Accuracy (ACC) and Chance level (CL) for Alpha=0.05; No days have a performance significantly (p>0.05) above chance level (not corrected for multiple comparisons).

| Day | D1 | D2 | D3 | D4 | D5 | D6 | D7 | D8 | D9 |
|---|---|---|---|---|---|---|---|---|---|
| Acc | 0.5813 | 0.46 | 0.4845 | 0.3523 | 0.5056 | 0.4133 | 0.5544 | 0.485 | 0.415 |
| CL | 0.6367 | 0.6674 | 0.5679 | 0.6184 | 0.564 | 0.6025 | 0.59 | 0.6025 | 0.6025 |

| Day | D10 | D11 | D12 | D13 | D14 | D15 | D16 | D17 | D18 |
|---|---|---|---|---|---|---|---|---|---|
| Acc | 0.53 | 0.4383 | 0.5475 | 0.4725 | 0.4725 | 0.4825 | 0.4525 | 0.425 | 0.5775 |
| CL | 0.6025 | 0.6025 | 0.5895 | 0.6236 | 0.6236 | 0.6236 | 0.5895 | 0.6025 | 0.6236 |

**Table A3.** Day-wise cross-validation results for Patient B with Accuracy (ACC) and Chance level (CL) for Alpha=0.05; No days have a performance significantly (p>0.05) above chance level (not corrected for multiple comparisons).

| Day | D1 | D2 | D3 | D4 | D5 | D6 | D7 | D8 | D9 | D10 | D11 | D12 |
|---|---|---|---|---|---|---|---|---|---|---|---|---|
| Acc | 0.54 | 0.5563 | 0.425 | 0.4788 | 0.416 | 0.518 | 0.517 | 0.4383 | 0.561 | 0.536 | 0.438 | 0.518 |
| CL | 0.5895 | 0.5895 | 0.6236 | 0.5895 | 0.589 | 0.602 | 0.589 | 0.6025 | 0.589 | 0.589 | 0.589 | 0.589 |

**Table A4.** Day-wise cross-validation results for Patient W with Accuracy (ACC) and Chance level (CL) for Alpha=0.05; No days have a performance significantly (p>0.05) above chance level (not corrected for multiple comparisons).

| Day | D1 | D2 | D3 | D4 | D5 | D6 |
|---|---|---|---|---|---|---|
| Acc | 0.57 | 0.475 | 0.50 | 0.53 | 0.55 | 0.55 |
| CL | 0.6025 | 0.6236 | 0.5895 | 0.5736 | 0.6025 | 0.6236 |

### A4. Statistical analysis of online classification results presented in (Guger et al., 2017)

In (Guger et al., 2017), a vibrotactile P300 BCI system is evaluated online in 9 patients (7 LIS, 2 CLIS). As the accuracy for all patients was at least 70 %, Guger et al. argue that the system worked for all patients. However, only a very small dataset (10 trials per patient) were collected and the online results were not evaluated statistically in that publication.

The statistical significance of a classification can be modeled by a binomial cumulative distribution (Combrisson et al., 2015). For n=10 trials, c=2 classes and a significance level $\alpha = 0.05$, the classification accuracy has to be greater (not equal to) 80 % to be significant. With the given number of trials, the actual p-value for 80 % is p=0.0546 and for 90 % p=0.0107.
Thereby, only 3 of the 9 online sessions (P1, P5, P6) have a classification accuracy with $p<0.05$. As significance was assessed individually for each patient, and 9 tests were performed, one would need to correct the p-values for multiple comparisons. When using a Bonferroni correction to correct for multiple comparisons, none of the patients has an online classification accuracy which is significantly above chance level.

With the online results not being significantly above chance level, the results presented in (Guger et al., 2017) should not be used to claim that the presented EEG-based BCI system established communication in the complete locked-in state.
However, the results warrant a further investigation, so Guger and colleagues are encouraged to test their system with a larger sample size (more trials per patient) and assess statistically if classification accuracy is significantly above chance level.

### A5. Inconsistencies and missing data in the supporting information to (Chaudhary et al., 2017)

For the analysis in this paper, all data published on Zenodo as supporting information to (Chaudhary et al., 2017) was used. As *Plos Biology* requires authors to make all data publicly available, Chaudhary et al. write in their Data Availability Statement that all data pertaining to the results in their paper can be found on Zenodo. Despite the statement, there are several inconsistencies and missing data when comparing the data published on Zenodo and the results presented in (Chaudhary et al., 2017). Due to data missing in the files uploaded to Zenodo, the number of sessions presented in this paper differs from the number of sessions presented in (Chaudhary et al., 2017) and not all aspects of the original paper could be reanalyzed.
In the following, all data pertaining to the results presented in (Chaudhary et al., 2017), but missing in the files published on Zenodo, are listed:

- raw fNIRS data for 30 training/feedback sessions for Patient F
- raw EEG and EOG data for all patients
- all data (EEG, EOG, fNIRS, labels) from open question sessions
- trained SVM models that were used in the feedback and open question sessions
- output of the online SVM classification for each trial in the feedback/open question sessions
