## Supplementary material for "No Evidence for Communication in the Complete Locked-in State": S2 Matlab Scripts used for analysis: analysis1.m

```
% This script was used for the producing the results of [1], which is a
% reanalysis of the data from [2].
% The data of [2] can be directly downloaded from:
% https://doi.org/10.5281/zenodo.192386 https://doi.org/10.5281/zenodo.192388
% https://doi.org/10.5281/zenodo.192390 https://doi.org/10.5281/zenodo.192391
%
% This scripts uses the LibSVM library for the SVM classification.
% LibSVM can be downloaded at: https://www.csie.ntu.edu.tw/~cjlin/libsvm/
%
% References:
% [1] M. Spüler (2018) Questioning the evidence for BCI-based communication in the complete locked-in state, submitted to Plos Biology
% [2] Chaudhary, U., Xia, B., Silvoni, S., Cohen, L. G., & Birbaumer, N. (2017). Brain–computer interface–based communication in the completely locked-in state. PLoS biology, 15(1), e1002593.


% load data

%fd=load('Fig2A+Table1_PatientF.mat')
%fd=load('Fig2B+Table1_PatientG.mat')
%fd=load('Fig2C+Table1_PatientB.mat')
fd=load('Fig2D+Table1_PatientW.mat')


data_x=[];
data_y=[];
data_sess=[];
data_day=[];

lastday='';
daycounter=0;
%put together all data
for sess=1:length(fd.tmp)
    sess_x=[];
    sess_y=[];
    sess_x(fd.ti(sess).trueIdx,:,:)= shiftdim(fd.HbXtrueNIRSdata(sess).data,2);
    sess_y(fd.ti(sess).trueIdx)=8;
    sess_x(fd.fi(sess).falseIdx,:,:)= shiftdim(fd.HbXfalseNIRSdata(sess).data,2);
    sess_y(fd.fi(sess).falseIdx)=4;
    sess_y=sess_y';
    newpos=size(data_x,1)+[1:size(sess_x,1)];
    data_x(newpos,:,:)=sess_x;
    data_y(newpos)=sess_y;
    data_sess(newpos)=sess;
    sss = strsplit([fd.datafile{sess}],'\');
    if (strcmp(sss{8}(1:10),lastday)~=1)
        lastday=sss{8}(1:10);
        daycounter=daycounter+1;        
    end
    data_day(newpos)=daycounter;        
     
end
% data_x contains HbX values with dimensions: trials X channel X samples
% data_y containes the label (4 no, 8 yes)
% data_sess is the session the individual trial belongs to


%% statistics
m4=squeeze(mean(data_x(data_y==4,:,:)));
m8=squeeze(mean(data_x(data_y==8,:,:)));

for ch=1:size(data_x,2)
    for smpl=1:size(data_x,3)
%         [h,p]=ttest(data_x(data_y==4,ch,smpl),data_x(data_y==8,ch,smpl));
%         rs(ch,smpl)=p;
       rs(ch,smpl) =ranksum(data_x(data_y==4,ch,smpl),data_x(data_y==8,ch,smpl));
    end
end
imagesc(rs<0.05)
title(['min p-value: ' num2str(min(min(rs)))])

%% Cross-validation
settings.cv_reps=1;
settings.numfolds=10;
settings.cv_stratified=1;
settings.opt_params=1; %optimize parameters: 1 yes, 0 no
settings.useslope=0;   % use slope 1 yes, 0 no (use raw data, which already represents the relative change)


cvacc=[];
res=[];

% calculate slope
data_slope=[];
if (settings.useslope==1)
    for trial=1:size(data_x,1)
        slope=data_x(trial,:,7:end)-data_x(trial,:,1:end-6);
        slope1s=slope(1,:,1:6:end);
        data_slope(trial,:,:)=slope1s;
    end
    data_x=data_slope;
end


for di=1:max(data_day) % do cross-validation for each day
    dx=data_x(data_day==di,:);
    dy=data_y(data_day==di)';
    
    for cvrep=1:settings.cv_reps % optional, repeat cross-validation multiple times for more stable results
        data_pred=[];
        inds_all=1:size(dx,1);
        inds_all = inds_all(randperm(length(inds_all)));
        for fold=1:settings.numfolds % cross-validation
            trials_test = inds_all(fold:settings.numfolds:end);
            trials_train = inds_all(~ismember(inds_all,trials_test));
            
            if (settings.cv_stratified) % stratified CV: balacing classes in training set
                minority_classamount = min(sum(dy(trials_train)==4),sum(dy(trials_train)==8));
                a= find(dy(trials_train)==4); a= a(randperm(length(a)));
                b= find(dy(trials_train)==8); b= b(randperm(length(b)));
                trials_train = trials_train([a(1:minority_classamount); b(1:minority_classamount)]);
            end
            
            %parameter optimization
            if settings.opt_params==1 
                testc=10.^[-8:8];
                for ti=1:length(testc)
                    svmparams=['-c ' num2str(testc(ti)) ' -s 0 -t 0 -v 10'];            
                    tacc(ti) = svmtrain(dy(trials_train),dx(trials_train,:),svmparams);
                end
                [a b]=max(tacc);
                usec=testc(b);
                
            else
                usec=1;
            end
            svmparams=['-c ' num2str(usec) ' -s 0 -t 0'];
            
            model = svmtrain(dy(trials_train),dx(trials_train,:),svmparams);
            [predicted_label, accuracy, decision_values]=svmpredict(dy(trials_test),dx(trials_test,:),model);
            data_pred(trials_test)=predicted_label;
        end
        acc(cvrep)=mean(data_pred'==dy);
        res(di).acc=mean(acc);
        n=length(dy);
        % get significance level threshold according to
        % Müller-Putz, G., Scherer, R., Brunner, C., Leeb, R., & Pfurtscheller, G. (2008). Better than random: a closer look on BCI results. International Journal of Bioelectromagnetism, 10, 52-55.
        p=(n*0.5+2)/(n+4);alpha=0.05;sig=p+sqrt(((p*(1-p))/(n+4)))*1.64;
        res(di).ci=sig;
    end  
end

Pw_res=[res.acc; res.ci;[res.acc]>[res.ci]];
```
