## Supplementary material for "No Evidence for Communication in the Complete Locked-in State": S2 Matlab Scripts used for analysis: analysis2_Patb.m

```
% This script was used for the producing the results of [1], which is a
% reanalysis of the data from [2].
% The data of [2] can be directly downloaded from:
% https://doi.org/10.5281/zenodo.192386 https://doi.org/10.5281/zenodo.192388
% https://doi.org/10.5281/zenodo.192390 https://doi.org/10.5281/zenodo.192391
%
%
% References:
% [1] M. Spüler (2018) No evidence for communication in the complete locked-in state, submitted to Plos Biology
% [2] Chaudhary, U., Xia, B., Silvoni, S., Cohen, L. G., & Birbaumer, N. (2017). Brain–computer interface–based communication in the completely locked-in state. PLoS biology, 15(1), e1002593.


fd=load('Fig2C+Table1_PatientB.mat')


%% Chaudhary et al. analysis: average first trials, then sessions, then channels
% used to produce Figure 1.A
data_x=[];
meant=[];
meanf=[];

for sess=1:length(fd.HbXtrueNIRSdata)
    tmp= cat(3,fd.HbXtrueNIRSdata(sess).data, fd.HbXfalseNIRSdata(sess).data);
    dt = tmp(:,:,[1:size(fd.HbXtrueNIRSdata(sess).data,3)]);
    df = tmp(:,:,[size(fd.HbXtrueNIRSdata(sess).data,3)+1:(size(fd.HbXtrueNIRSdata(sess).data,3)+size(fd.HbXfalseNIRSdata(sess).data,3))]);
    meant(sess,:,:)=squeeze(mean(dt,3));
    meanf(sess,:,:)=squeeze(mean(df,3));
end
meants=squeeze(mean(meant,1));
meanfs=squeeze(mean(meanf,1));
[tth ttp]=(ttest(meants,meanfs));
figure;
plotmeans(meants,meanfs,ttp<0.0005)


%% Chaudhary et al. analysis with randomly permuted trials
% used to produce Figure A1 in Appendix
data_x=[];
meant=[];
meanf=[];

figure;
for p=1:10
    rng(p);
    for sess=1:length(fd.HbXtrueNIRSdata)
        tmp= cat(3,fd.HbXtrueNIRSdata(sess).data, fd.HbXfalseNIRSdata(sess).data);
        tmp = tmp(:,:,randperm(size(tmp,3)));
        dt = tmp(:,:,[1:size(fd.HbXtrueNIRSdata(sess).data,3)]);
        df = tmp(:,:,[size(fd.HbXtrueNIRSdata(sess).data,3)+1:(size(fd.HbXtrueNIRSdata(sess).data,3)+size(fd.HbXfalseNIRSdata(sess).data,3))]);
        meant(sess,:,:)=squeeze(mean(dt,3));
        meanf(sess,:,:)=squeeze(mean(df,3));
    end
    meants=squeeze(mean(meant,1));
    meanfs=squeeze(mean(meanf,1));
    [tth ttp]=(ttest(meants,meanfs));

    subplot(2,5,p);
    plotmeans(meants,meanfs,ttp<0.0005)
end

%% corrected analysis: average first channels, then trials, then sessions
% used to produce Figure 1.B
data_x=[];
meant=[];
meanf=[];
for sess=1:length(fd.HbXtrueNIRSdata)
    tmp= cat(3,fd.HbXtrueNIRSdata(sess).data, fd.HbXfalseNIRSdata(sess).data);
    dt = tmp(:,:,[1:size(fd.HbXtrueNIRSdata(sess).data,3)]);
    df = tmp(:,:,[size(fd.HbXtrueNIRSdata(sess).data,3)+1:(size(fd.HbXtrueNIRSdata(sess).data,3)+size(fd.HbXfalseNIRSdata(sess).data,3))]);
    
    meants(sess,:)=squeeze(mean(mean(dt,1),3));
    meanfs(sess,:)=squeeze(mean(mean(df,1),3));
end

[tth ttp]=(ttest(meants,meanfs));
figure;
plotmeans(meants,meanfs,ttp<0.05)
```
