## Supplementary material for "No Evidence for Communication in the Complete Locked-in State": S2 Matlab Scripts used for analysis: plotmeans.m

```
function plotmeans(meants,meanfs,tth)
    
    h=plot(mean(meants,1),'r');hold on;    
    set(h,'LineWidth',2)
    
    h=plot(mean(meanfs,1),'b');
    set(h,'LineWidth',2)
    
    x=1:size(meanfs,2);
    y1=mean(meants,1)-std(meants,1);
    y2=mean(meants,1)+std(meants,1);
    h=fill([x x(end:-1:1)], [y1 y2(end:-1:1)],'r');
    set(h,'facealpha',.5);
    set(h,'LineStyle','none');


    x=1:size(meanfs,2);
    y1=mean(meanfs,1)-std(meanfs,1);
    y2=mean(meanfs,1)+std(meanfs,1);
    h=fill([x x(end:-1:1)], [y1 y2(end:-1:1)],'b');
    set(h,'facealpha',.5);
    set(h,'LineStyle','none');


    ticks=get(gca,'YTick');

    startsig=-1;
    endsig=-1;
    for i=1:length(meants)
        if (tth(i)==1 && startsig<=0)
            startsig=i;
        end
        if ( (tth(i)==0 || i==length(meants)) && startsig>0)
            endsig=i-1;
            if (tth(i)==1) %check for last sample
                endsig=i;
            end
            h=fill([startsig-0.5, endsig+0.5, endsig+0.5, startsig-0.5],[ticks(1), ticks(1), ticks(end), ticks(end)],[0.5 0.5 0.5]);
            set(h,'facealpha',.2);
            set(h,'LineStyle','none');

            startsig=-1;
            endsig=-1;
        end
    end
    xlim([1 length(meants)]);
```
