## Supplementary material for "No Evidence for Communication in the Complete Locked-in State": S2 Matlab Scripts used for analysis: readme_scripts.txt

analysis1.m  
This script was used to generate the first statistical analysis (shown in Appendix), as well as the classification results

analysis2_Patb.m 
This script was used for the second statistical analysis and to plot the figures in the manuscript.

plotmeans.m
An additional script needed for plotting
